# Infertility due to defective sperm flagella caused by an intronic deletion in DNAH17 that perturbs splicing

**DOI:** 10.1101/2020.10.16.333344

**Authors:** Adéla Nosková, Maya Hiltpold, Fredi Janett, Thomas Echtermann, Zih-Hua Fang, Xaver Sidler, Christin Selige, Andreas Hofer, Stefan Neuenschwander, Hubert Pausch

## Abstract

Artificial insemination in pig (*Sus scrofa domesticus*) breeding involves the evaluation of the semen quality of breeding boars. Ejaculates that fulfill predefined quality requirements are processed, diluted and used for inseminations. Within short time, eight Swiss Large White boars producing immotile sperm that had multiple morphological abnormalities of the sperm flagella were noticed at a semen collection center. The eight boars were inbred on a common ancestor suggesting that the novel sperm flagella defect is a recessive trait. Transmission electron microscopy cross-sections revealed that the immotile sperm had disorganized flagellar axonemes. Haplotype-based association testing involving microarray-derived genotypes at 41,094 SNPs of six affected and 100 fertile boars yielded strong association (P=4.22 × 10^−15^) at chromosome 12. Autozygosity mapping enabled us to pinpoint the causal mutation on a 1.11 Mb haplotype located between 3,473,632 and 4,587,759 bp. The haplotype carries an intronic 13-bp deletion (Chr12:3,556,401-3,556,414 bp) that is compatible with recessive inheritance. The 13-bp deletion excises the polypyrimidine tract upstream exon 56 of *DNAH17* (XM_021066525.1:c.8510-17_8510-5del) encoding dynein axonemal heavy chain 17. Transcriptome analysis of the testis of two affected boars revealed that the loss of the polypyrimidine tract causes exon skipping which results in the in-frame loss of 89 amino acids from DNAH17. Disruption of DNAH17 impairs the assembly of the flagellar axoneme and manifests in multiple morphological abnormalities of the sperm flagella. Direct gene testing may now be implemented to monitor the defective allele in the Swiss Large White population and prevent the frequent manifestation of a sterilizing sperm tail disorder in breeding boars.

## INTRODUCTION

Artificial insemination is the most frequent method of breeding in pigs. The semen of breeding boars is collected once or twice per week at semen collection centers.

Traits that are routinely measured in all ejaculates include ejaculate volume, sperm concentration, motility, and morphology. Only ejaculates that meet predefined quality requirements are processed, diluted and used for inseminations (Colenbrander *et al.*, 1993),(Holt *et al.*, 1997),(Broekhuijse *et al.*, 2011). Semen quality and insemination success vary within and between boars due to environmental, permanent environmental and genetic effects (Marques *et al.*, 2017).

Access to longitudinal data on standardized semen traits for large cohorts of males is unique to livestock populations (Thibier and Wagner, 2002),(Knox, 2016). Boars suitable for breeding are selected based on semen quality and genomic predictions derived from dense SNP microarray-derived genotypes. Dense genotypes and repeated semen trait measurements for thousands of males are prerequisites to unravel the genetic architecture of male fertility. The comprehensive genetic analysis of male fertility is challenging in most species other than livestock. For instance, the analysis of male fertility in humans typically relies on correlated proxy phenotypes such as family size and birth rate (Kosova *et al.*, 2012), small cohorts of males (Sato *et al.*, 2018),(Sato *et al.*, 2020), and observations from a single ejaculate per individual (Rahban *et al.*, 2019).

Boar semen quality is positively correlated with insemination success and litter size (Holt *et al.*, 1997). Semen traits that are measured at semen collection centers have medium heritability (Marques *et al.*, 2017). Thus, they are well suited response variables in genome-wide association studies for male reproductive performance (Diniz *et al.*, 2014),(Hiltpold *et al.*, 2020). Genome-wide association studies on male fertility carried out in livestock discovered novel phenotype-genotype associations that improved our biological understanding of mammalian fertilization (Pausch *et al.*, 2014),(Lamas-Toranzo *et al.*, 2020),(Noda *et al.*, 2020),(Hiltpold *et al.*, 2020),(Ni *et al.*, 2020),(Noskova *et al.*, 2020).

Ejaculates from hundreds of boars are evaluated every year at semen collection centers as a service to the pig breeding industry. This rich resource of semen quality records facilitates to investigate sporadically occurring sperm defects using case-control association testing (Sironen *et al.*, 2006),(Sironen *et al.*, 2011),*(Noskova et al.*, 2020). Once causative variants have been identified, direct gene tests and genome-based mating strategies may be implemented to avoid the birth of infertile males. Moreover, the discovery of genes that harbor pathogenic alleles that compromise male fertility is important to enhance the diagnostic yield of genetic testing also in species other than livestock (Xavier *et al*., 2020).

Here, we investigate an autosomal recessive sperm tail defect of Swiss Large White boars. Using genome-wide association testing, we map the disorder to porcine chromosome 12. The analysis of genome-wide DNA and RNA sequencing data of affected boars revealed that the loss of an intronic polypyrimidine tract of porcine *DNAH17* is causal for the morphological abnormalities of the sperm flagella.

## MATERIAL AND METHODS

### Ethics approval and consent to participate

Semen samples of five breeding boars were collected by laboratory technicians at an approved semen collection center as part of their regular service to the Swiss pig breeding industry. Semen samples of three boars were flushed from the epididymis post mortem. Testes were collected after regular slaughter at an approved slaughterhouse. Our study was approved by the veterinary office of the Canton of Zurich (animal experimentation permit ZH 070/20).

### Consent for publication

SUISAG, the Swiss pig breeding and competence center provided written consent to the analyses performed and agreed to publish results and data.

### Animals

Eight Swiss Large White boars with a sperm tail defect were considered in our study (**Table 1**). Five of them were noticed at the semen collection center of SUISAG because their ejaculates contained immotile spermatozoa that had multiple morphological abnormalities of the flagella. Because the five boars were healthy and pedigree analysis indicated that they were inbred on a common ancestor, recessive inheritance of the sperm tail defect was suspected. A mating between two suspected carrier animals was performed in the field to assess phenotypic manifestations in their offspring. The pregnant sow was purchased and maintained at the research barn of the Division of Swine Medicine, Vetsuisse Faculty, University of Zurich. The sow gave birth to a litter with eleven piglets (eight females, three males). One of the male piglets (Boar_1246) died at the age of 200 days due to hemorrhagic bowel syndrome (Grahofer *et al.*, 2017). The other two male boars (Boar_1249, Boar_1254) were slaughtered at the age of 17 months at a regular slaughterhouse. All three male piglets from the mating of suspected carrier animals expressed the sperm tail defect (see below). Pedigree records were analyzed using the PYPEDAL software package (Cole, 2007).

**Table 1:**
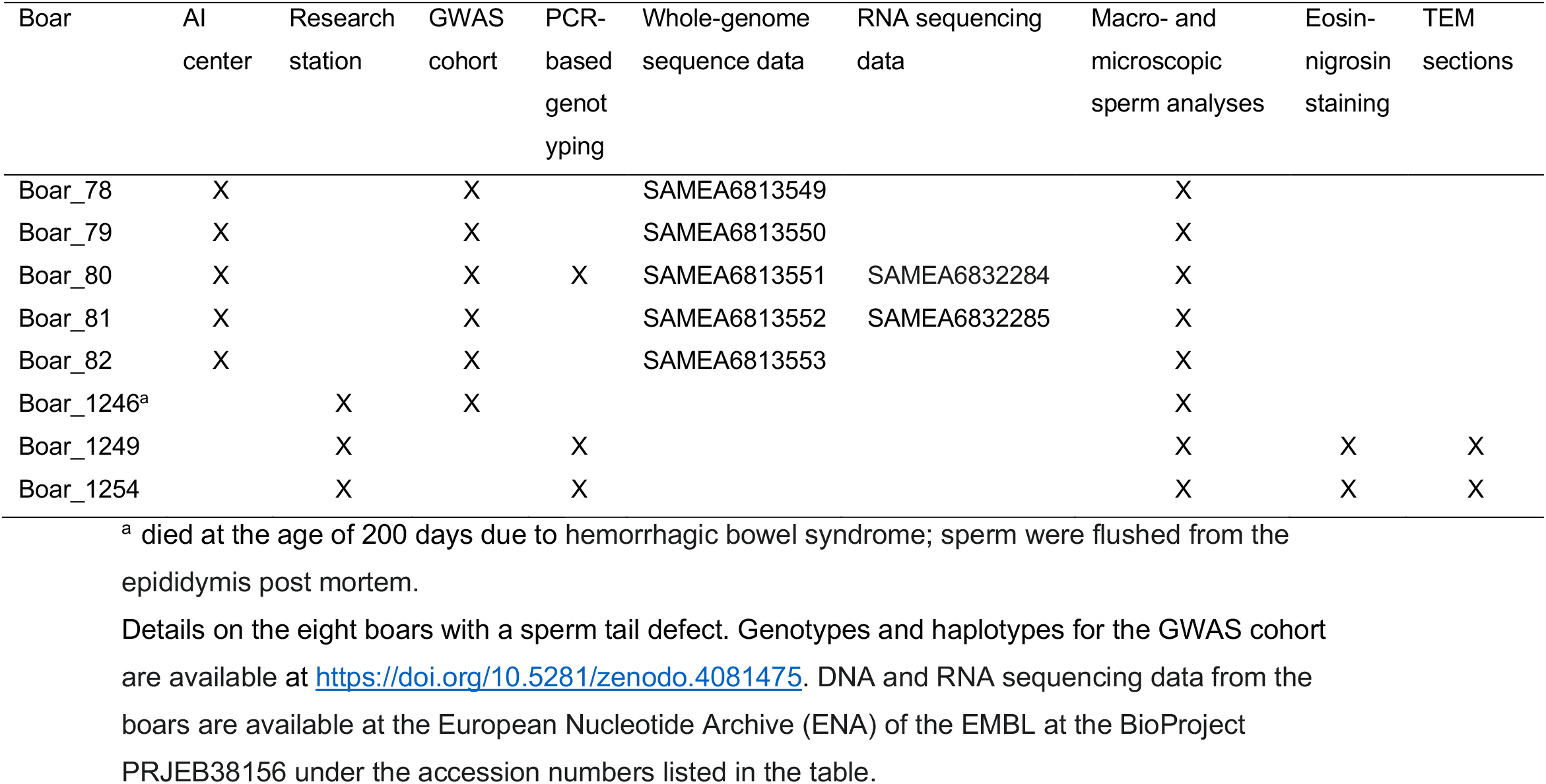
Experimental design of the study.

### Phenotypes

Semen samples of five breeding boars were macro- and microscopically evaluated by laboratory technicians as part of the routine service of SUISAG to the Swiss pig breeding industry. Spermatozoa from three boars (Boar_1246, Boar_1249, Boar_1254) were flushed from the epididymis post mortem. Semen samples from a fertile boar were provided by SUISAG.

Sperm concentration, total sperm count and sperm motility were determined with an IVOS II CASA system (Hamilton Thorne Inc., Beverly, U.S.A.) using Leja 2-chamber slides (Leja, Nieuw-Vennep, the Netherlands). For morphological examination, semen was fixed in buffered formol saline solution (Na2HPO4 4.93g, KH2PO4 2.54g, 38% formaldehyde 125ml, NaCl 5.41g, distilled water q.s. 1000ml) and smears prepared (Hancock, 1957). At least 200 spermatozoa were subsequently evaluated by phase contrast microscopy using oil immersion (Olympus BX50, UplanF1 100×/1.30, Olympus, Wallisellen, Switzerland).

Assessment of sperm viability was performed using the eosin-nigrosin staining method (Blom, 1950). In stained slides at least 200 spermatozoa were evaluated under oil immersion on a light microscope (Olympus BX50, UplanF1 100×/1.30, Olympus, Wallisellen).

In order to prepare sperm flushed from the epididymis for transmission electron microscopy (TEM), the samples were washed twice in phosphate buffered saline (PBS) and subsequently centrifuged at 300g. A semen sample from an unaffected boar was first centrifuged at 300g to increase the sperm concentration and then washed in PBS. The pellet was fixed with equal volume of 6% glutaraldehyde, resuspended gently, and centrifuged at 6000g. After removing the supernatant, the sperm were fixed for a second time with 3% glutaraldehyde, and finally pelleted at 6000g. Pellets were washed three times in PBS, post-fixed in 1% osmium tetroxide, washed in ddH2O, stained in 1% uranyl acetate, dehydrated in graded series of ethanol (25, 50, 75, 90, and 100%), and embedded in Epoxy resin through increasing concentrations (25, 50, 75, and 100%) using PELCO Biowave+ tissue processor, and then cured at 60 °C for 3 days. Embedded blocks were sectioned using Leica FC6 microtome and a DIATOME diamond knife with 45° angle into 60 nm sections and mounted on Quantifoil copper grids with formvar carbon films. Sections were post-stained with 2% uranyl acetate followed by lead citrate. Grids were imaged using FEI Morgagni 268 electron microscope operated at 100 kV at 20 k magnification.

### Genotypes and haplotype inference

Genotypes of 9955 Large White boars (including six boars with the sperm tail defect) and sows were provided by SUISAG. All pigs were genotyped using Illumina PorcineSNP60 Bead chips that comprised between 62,163 and 68,528 SNPs. We considered only autosomal SNPs. Physical positions of the SNPs corresponded to the Sscrofa11.1-assembly of the porcine genome (Warr *et al.*, 2019). Quality control on the genotypes was carried using the PLINK (version 1.9) software (Chang *et al.*, 2015). Animals and SNPs with more than 10% missing genotypes were excluded from subsequent analyses. We removed SNPs with minor allele frequency (MAF) less than 0.005 and SNPs for which the observed genotype distribution deviated significantly (P < 0.00001) from Hardy-Weinberg proportions. After quality control, our dataset contained 9848 pigs and genotypes at 43,254 autosomal SNPs. Sporadically missing genotypes were imputed and haplotypes were inferred using 12 iterations of the phasing algorithm implemented in the BEAGLE (version 5.0) software (Browning *et al.*, 2018), assuming an effective population size of 500 while all other parameters were set to default values.

### Genome-wide association testing

Six boars that produced sperm with multiple morphological abnormalities of the flagella were considered as case group for a genome-wide association study. The control group consisted of 100 randomly selected boars that produced normal sperm and were fertile, i.e., each of them sired at least one litter in the Swiss Large White breeding unit.

We tested the association between the affection status of the boars and 41,094 autosomal SNPs for which the frequency of the minor allele was greater than 5%. Single marker-based case-control association testing was performed using Fisher’s exact tests of allelic association as implemented in the PLINK (version 1.9) software (Chang *et al.*, 2015). In order to take population stratification into account, we performed also a SNP-based mixed linear model-based association study using the *mlma* method of the GCTA (version 1.92.1) software (Yang *et al.*, 2011).

A linear haplotype-based association model was implemented as described in (Venhoranta *et al*., 2014) and (Hiltpold *et al.*, 2020). We shifted a sliding window consisting of 25 contiguous SNPs corresponding to an average haplotype length of 1.28 ± 0.65 Mb along the chromosomes in steps of 5 SNPs. Within each of the 8583 sliding windows considered, we tested the association of between 1 and 10 haplotypes that had frequency greater than 5% with the sperm tail defect using the linear model 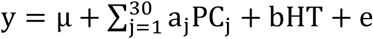, where y is a vector of phenotypes (1-fertile, 2-infertile), μ is the intercept, PC_j_ are the top 30 principal components of a genomic relationship matrix built based on genotypes of 43,254 autosomal SNPs using the PLINK software, a and b are effects of the principal components and the haplotype (HT) tested, respectively, and e is a vector of residuals that are assumed to be normally distributed. In total, we tested 47,685 haplotypes for association with the sperm tail defect.

SNPs and haplotypes that exceeded the Bonferroni-corrected significance threshold (P_SNP_=1.22 × 10^−6^, P_HAPLO_=1.05 × 10^−6^) were considered as significantly associated. The inflation factors of the test statistics were calculated using the *estlambda()*-function of the GENABEL R package (Aulchenko *et al.*, 2007).

### Analysis of positional candidate genes

The top association signal revealed a 1.11 Mb segment of extended homozygosity at chromosome 12. Genes located within the segment of homozygosity were obtained from the Refseq annotation of the porcine genome (version 106, available at ftp://ftp.ncbi.nlm.nih.gov/genomes/refseq/vertebrate_mammalian/Sus_scrofa/annotation_releases/106/GCF_000003025.6_Sscrofa11.1/GCF_000003025.6_Sscrofa11.1_genomic.gff.gz). Using publicly available RNA sequencing data (BioProject PRJNA506525, sample accession numbers SAMN10462191 – SAMN10462197) from (Robic *et al.*, 2019), we quantified mRNA abundance of the positional candidate genes in porcine testes. To this end, we downloaded between 47 and 58 million paired-end (2 × 100 and 2 × 125 bp) sequencing reads that were generated from RNA extracted from testis tissue of seven pubertal boars from European Piétrain and Piétrain × Large White populations. The RNA sequencing reads were pseudo-aligned to an index of the porcine transcriptome (Refseq version 106) and transcript abundance was quantified using the KALLISTO software (Bray *et al.*, 2016). We used the R package TXIMPORT (Soneson *et al.*, 2015) to aggregate transcript abundances to the gene level.

### Whole-genome sequencing and sequence variant genotyping

We sequenced five Swiss Large White boars that produced sperm that had multiple morphological abnormalities of the sperm flagella using 2 ×150 bp paired-end reads. Illumina TruSeq DNA PCR-free libraries with 400 bp insert sizes were prepared and sequenced at an Illumina NovaSeq6000 instrument. We used the FASTP software (Chen *et al.*, 2018) to remove adapter sequences, poly-G tails and reads that had phred-scaled quality less than 15 for more than 15% of the bases. Between 81,145,807 and 114,616,836 filtered read pairs were aligned to the Sscrofa11.1 assembly of the porcine genome using the MEM algorithm of the BWA software (Li, 2013) with default parameter settings. We marked duplicates using the PICARD tools software suite (https://github.com/broadinstitute/picard) and sorted the alignments by coordinates using SAMBAMBA (Tarasov *et al.*, 2015). The number of reads covering a position were extracted from the alignments using the MOSDEPTH software (version 0.2.2) with default parameter settings (Pedersen and Quinlan, 2018). The realized sequencing read depth of the five boars was 10.9-fold and it ranged from 8.9-to 12.5-fold.

Sequence variants (SNPs and Indels) of the five sequenced boars were genotyped together with 93 pigs from different breeds for which whole-genome sequence data were available from our in-house database using the multi-sample variant calling approach of the GENOME ANALYSIS TOOLKIT (GATK, version 4.1.0; (Depristo *et al.*, 2011)). We filtered the sequence variants by hard-filtering according to best practice guidelines of the GATK. A detailed description of our reference-guided variant discovery and filtration approach is described in (Crysnanto *et al.*, 2019). Functional consequences of polymorphic sites were predicted according to the Refseq annotation (version 106) of the porcine genome using the VARIANT EFFECT PREDICTOR software from Ensembl (McLaren *et al.*, 2016) along with the SpliceRegion.pm plugin (https://github.com/Ensembl/VEP_plugins/blob/release/101/SpliceRegion.pm). We considered 82 pigs with known pedigree as a control group for the present study.

### Identification of candidate causal variants

We considered 14,806 SNPs and 3,856 Indels that were detected within the 1.11 Mb segment (between 3,473,632 and 4,587,759 bp) of extended homozygosity at chromosome 12 as positional candidate causal variants. To identify variants compatible with recessive inheritance of the sperm tail disorder, we applied a filtering strategy that takes into account flawed genotypes and the under-calling of heterozygous genotypes due to relatively low sequencing coverage. Specifically, we screened for alleles that had following frequency:

- ≤ 0.8 in five affected boars (at least 8 out of 10 alleles),
- ≥ 0.05 in 82 control pigs from different breeds (less than 8 alleles).

This filtration resulted in only one compatible variant which was a 13-bp intronic deletion.

### Whole transcriptome sequencing and read alignment

Testes were collected immediately after slaughter at an approved slaughterhouse from two boars (Boar_80, Boar_81) that were homozygous for the 13-bp deletion and produced sperm with multiple morphological abnormalities of the flagella. Tissue samples were frozen in liquid nitrogen and stored at −80°C until RNA extraction. Paired-end RNA libraries (2 × 150 bp) were prepared using total RNA purified from testis tissue using the Illumina TruSeq RNA Sample Preparation Kit (Illumina, San Diego, CA, USA). The libraries were sequenced at an Illumina NovaSeq6000 instrument yielding 49,198,010 and 70,451,148 reads. Quality control on the raw RNA sequencing reads was performed using the FASTP software using default parameter settings (see above). The filtered read pairs (48,889,296 and 69,988,376) were aligned to the Sscrofa11.1 reference sequence and the Refseq gene annotation (version 106) using the splice-aware read alignment tool STAR (version 2.7.3a) (Dobin *et al.*, 2013). The number of RNA sequencing reads that covered a position was counted using the MOSDEPTH (see above).

### Bioinformatic analysis

Putative branch point sequences within *DNAH17* (XM_021066525.1) intron 55 were predicted using the BPP algorithm (Zhang *et al.*, 2017). Putative 3’ splice acceptor sites were predicted using the NNSPLICE software tool (https://www.fruitfly.org/seq_tools/splice.html (Reese *et al.*, 1997)). Domains of porcine DNAH17 (protein-ID: A0A287AFU3) were retrieved from Uniprot.

### Genotyping of the 13-bp deletion

DNA was isolated from either hair roots (living pigs) or spleen (slaughtered pigs) using DNeasy blood and tissue kit (Qiagen). A PCR test was established consisting of a FAM-labelled forward primer (Fw-*DNAH17*: 5’-TGAGCATCTTCTTGGCGAGG-3’) and a reverse primer (Re-*DNAH17*: 5’-GCTGTTGATCAGCACCAGGA-3’) which bind to the wild type and mutant alleles, as well as a wild type specific reverse primer (Re-wt-*DNAH17*: 5’-TCGAGCTAGACGCGAGGG-3’). PCR fragments were analyzed on a DNA analyzer 3130xl (Applied Bioscience).

### Availability of data

Whole-genome sequence data of 87 pigs including five boars with a sperm tail defect have been deposited at the European Nucleotide Archive (ENA) of the EMBL at BioProject PRJEB38156, PRJEB37956, PRJEB39374 and PRJNA622908, under sample accession numbers listed in **Supplemental file S1**. The genotypes of five affected and 82 unaffected boars at 14,806 SNPs and 3,856 Indels that were detected within the segment of autozygosity are available from Zenodo (https://doi.org/10.5281/zenodo.4081475). RNA sequencing data of testicular tissue samples of two boars homozygous for the 13-bp deletion have been deposited at the European Nucleotide Archive (ENA) of the EMBL at the BioProject PRJEB38156 under sample accession numbers SAMEA6832284 and SAMEA6832285. RNA sequencing data of testicular tissue samples of seven pubertal boars are accessible via the European Nucleotide Archive (ENA) of the EMBL at the BioProject PRJNA506525 under sample accession numbers SAMN10462191 – SAMN10462197. The microarray-derived genotypes and haplotypes of cases and controls, the eigenvectors considered to account for stratification, the R script used to carry out haplotype-based association testing, and the results from the haplotype-based association study are available from Zenodo (https://doi.org/10.5281/zenodo.4081475).

## RESULTS

### Phenotypic manifestation of a sperm tail disorder in Large White boars

Five Swiss Large White boars maintained at a semen collection center produced ejaculates that contained spermatozoa with defective tails. Ejaculate volume and sperm concentration were normal. Microscopic semen analysis revealed multiple morphological abnormalities of the sperm flagella including rudimentary, short, coiled and irregularly shaped tails (**Figure 1A**). Proximal cytoplasmic droplets were frequently observed at the junction of sperm head and tail. Progressively motile sperm were not detected in the ejaculates. Eosin-nigrosin staining of semen flushed from the epididymis of two affected boars indicated that 48 and 60% of the sperm were viable (**Supplemental file S2**). Apart from producing sperm with defective tails, the boars were healthy and had no testicular abnormalities. Due to the absence of motile sperm and the high degree of abnormalities of the flagella, the boars were not suitable for breeding and were slaughtered. The sperm tail defect of the boars is commonly referred to as asthenoteratozoospermia.

**Figure 1.**
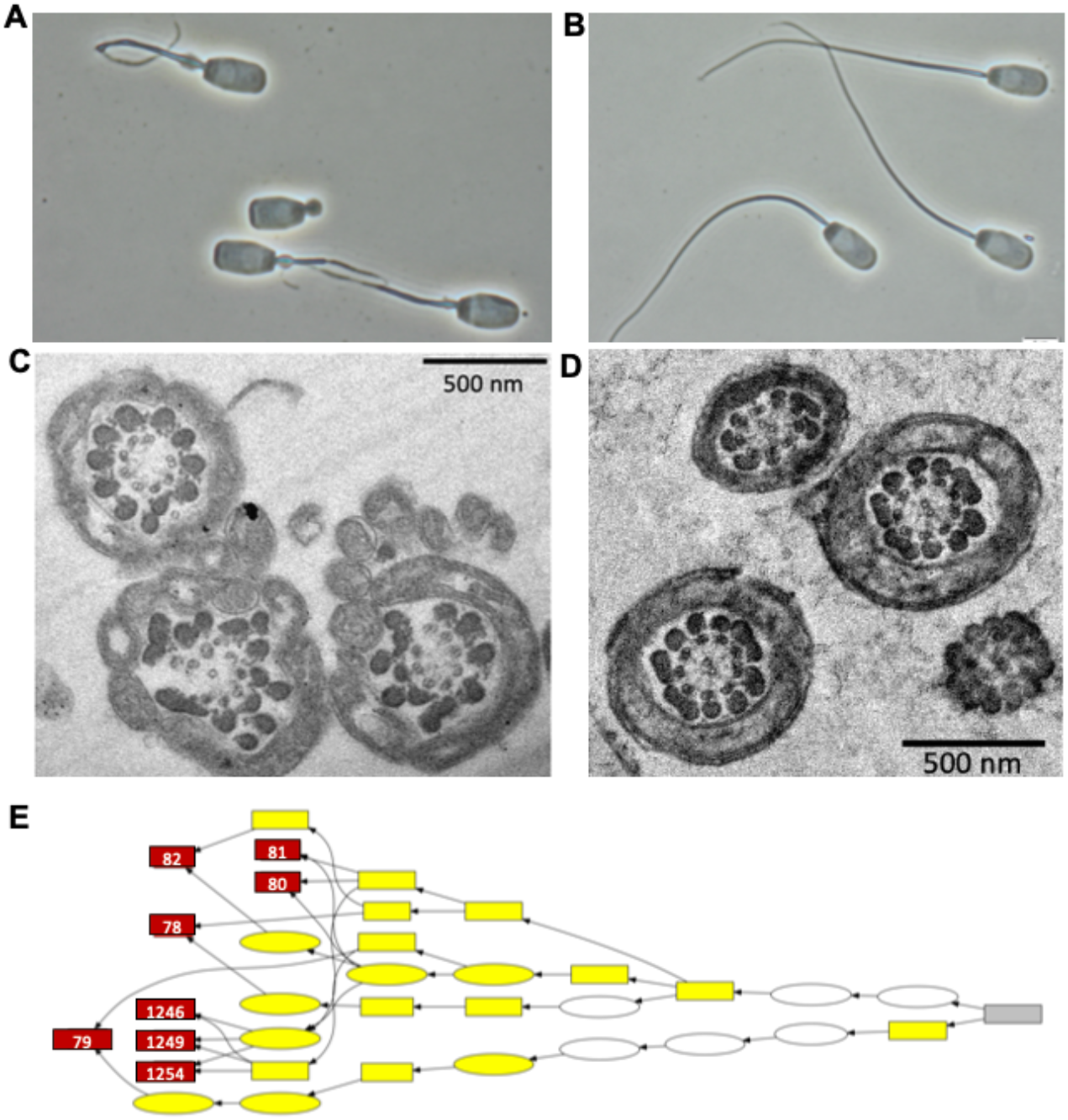
Characteristics of a recessive sperm tail defect of Large White boars. Representative phase contrast microscopy images of sperm produced from affected (A) and fertile (B) boars. Ejaculates of affected boars contained sperm with multiple morphological abnormalities of the flagella including short, coiled, broken and bent tails. A proximal cytoplasmic droplet was detected between the sperm head and tail at most of the spermatozoa from affected boars. Representative transmission electron microscopy cross-sections of sperm flagella from affected (C) and fertile (D) boars. Cross sections of the sperm flagella revealed the typical arrangement of nine outer microtubule doublets surrounding the central pair (9×2 + 2) in fertile boars (D). Lack of central and outer microtubules doublets and complete axonemal disorganization were frequently detected in cross-sections of the sperm flagella in affected boars. Pedigree of eight boars that produced sperm with defective tails (D). The pedigree contains only obligate mutation carriers. Red color indicates the eight affected boars. The numbers within the boxes correspond to the IDs in **Table 1**. Boars 1246, 1249 and 1254 originate from a mating between two suspected carrier animals. Ovals and boxes represent female and male ancestors, respectively. The grey box indicates a common ancestor (born in 2005) on which all affected boars were inbred. Shapes filled with yellow color represent individuals that carried the top haplotype in the heterozygous state (see below).

Transmission electron microscopy cross-sections of sperm flagella from a fertile boar revealed the typical axonemal arrangement of nine outer microtubule doublets surrounding the central pair (9×2 + 2) (**Figure 1D**). In contrast, cross-sections of sperm that were flushed from the epididymis of two affected boars revealed multiple ultrastructural abnormalities including completely disorganized axonemes (**Figure 1C**). Central and outer microtubule doublets were absent in some flagella. Some sperm had superfluous but disorganized axonemal structures. The cytoplasmic bags that were detected at the junction between sperm head and tail using light microscopy contained unassembled axonemal components or multiple flagellar-like structures within the cell membrane of one sperm (**Supplemental file S3**).

The boars that produced defective sperm were closely related (**Figure 1E**). Two affected boars were fullsibs (Boar_80, Boar_81). The average relationship coefficient between the boars was 0.28 and it ranged from 0.14 to 0.63. Their average coefficient of inbreeding was 0.087, which is slightly higher than in the fertile boars (F=0.06). All affected boars were inbred on a boar born in 2005. This common ancestor was present in both the paternal and maternal ancestry of significantly less (N=10, P_Fisher’s exact_=4.69 × 10^−6^) fertile boars. Three sires of affected boars were used in artificial insemination. None of their ejaculates (n=168) contained an anomalous amount of sperm with defective flagella. These observations are compatible with an autosomal recessive inheritance of the sperm tail defect.

In order to verify the presumed recessive inheritance and monitor phenotypic manifestations in homozygous boars, a mating between two suspected carrier animals was performed in the field (**Figure 1E**). The sow gave birth to a litter with eight female and three male piglets (Boar_1246, Boar_1249, Boar_1254). The three male piglets were maintained at a research barn. Boar_1246 died at the age of 200 days due to hemorrhagic bowel syndrome. Testes and epididymis were collected post mortem. Testicular abnormalities were not detected. Microscopic analysis of sperm flushed from the epididymis revealed that the sperm flagella had multiple morphological abnormalities. Boar_1249 and Boar_1254 were healthy and developed normal at the research barn. At 17 months, both boars were slaughtered at a regular slaughterhouse and testes and epididymis were collected. Microscopic analysis of sperm flushed from the epididymis revealed that both boars produced sperm with multiple morphological abnormalities of the flagella.

### Haplotype-based association testing maps the sperm tail defect to a 1.11 Mb interval on porcine chromosome 12

Six affected boars (**Table 1**) were genotyped with the Illumina PorcineSNP60 microarray. As a control group, we considered 100 randomly selected fertile boars of the Swiss Large White breeding population that also had Illumina PorcineSNP60-derived genotypes. Following quality control, genotypes at 41,094 autosomal SNPs were used for association testing. Single marker-based case/control-association testing using Fisher’s exact test of allelic association revealed eleven SNPs located on porcine chromosomes 11, 12 and 13 that exceeded the Bonferroni-corrected significance threshold (**Figure 2A**). The strongest association signal (P=1.87 × 10^−9^) resulted from SNP *ASGA0052524* that is located at 2,785,333 bp on porcine chromosome 12. An inflation factor of 1.55 indicated that the SNP-based association study was enriched for false positive association signals. Inspection of the top principal components of the genomic relationship matrix revealed clustering of the boars with the sperm tail defect (**Supplemental file S4**), suggesting that population stratification confounded the association analysis, thereby producing spurious association signals.

**Figure 2.**
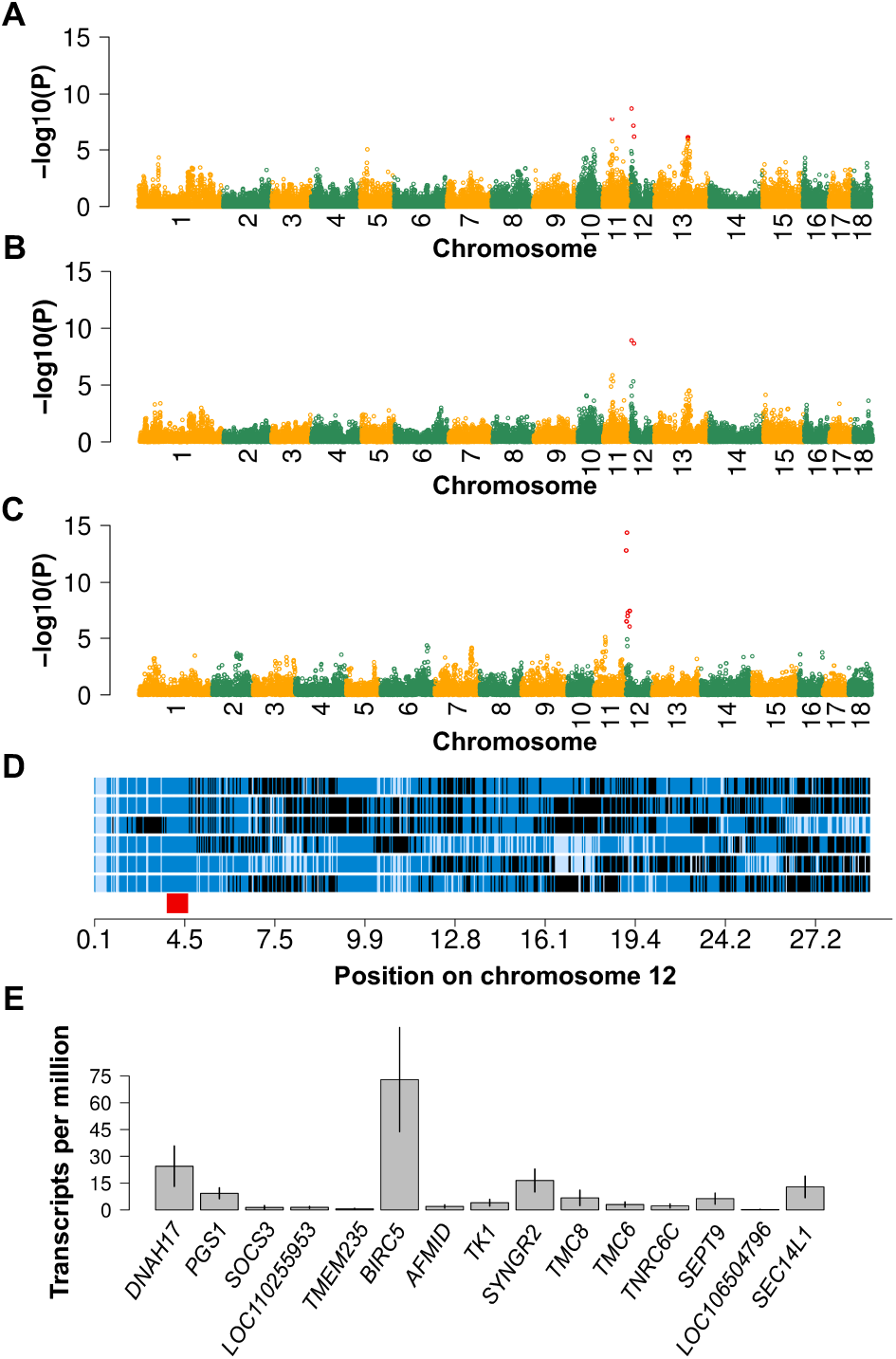
A 1.11 Mb segment on porcine chromosome 12 is associated with the sperm tail defect. Manhattan plot of a SNP-(A, B) and haplotype-based (C) genome-wide association study. Red color represents eleven SNPs and twenty-one haplotypes that exceeded the Bonferroni corrected-significance threshold. SNP-based association studies were performed using either Fisher exact tests of allelic association (A) or a mixed linear model as implemented in GCTA (B). Homozygosity mapping in six boars that produced sperm with defective tails using SNPs located on porcine chromosome 12 (D). Blue and pale blue represent homozygous genotypes (AA and BB), heterozygous genotypes are displayed in black. The red bar indicates a 1.11 Mb segment of extended homozygosity that was shared in all six affected boars. mRNA abundance (quantified in transcripts per million) of 15 genes that are located within the segment of extended homozygosity in testicular tissue of pubertal boars (E). The vertical line represents the standard deviation estimated across seven samples.

To take population stratification into account, we repeated the association analysis using either a SNP-based mixed linear model or a haplotype-based linear model that included 30 principal components of a genomic relationship matrix as covariates. An inflation factor of 1.03 indicated that the SNP-based mixed linear model successfully controlled population stratification. Two SNPs (ASGA0052524 at 2,785,333 bp and ALGA0118426 at 8,504,544 bp) at SSC12 were significantly associated with the sperm tail defect (**Figure 2B**).

An inflation factor of 1.15 indicated that the principal components-based control of population stratification was mostly successful in the haplotype-based association study. Twenty-one haplotypes located between 94,345 and 9,256,260 bp on chromosome 12 (SSC12) exceeded the Bonferroni-corrected significance threshold (**Figure 2C**). The strongest associations (P=4.22 × 10^−15^) were detected for three overlapping haplotypes located between 2,166,472 and 3,755,956 bp. The top haplotype occurred at a frequency of 3.5% among the 100 fertile boars. Seven fertile boars were heterozygous carriers of the top haplotype. The top haplotype was not detected in the homozygous state in fertile boars. No haplotypes located on chromosomes other than SSC12 were associated with the sperm tail defect.

The haplotype-based association study peaked at a 1.11 Mb segment of extended homozygosity (between 3,473,632 and 4,587,759 bp) at porcine chromosome 12 that was identical by descent in all six affected boars, corroborating recessive inheritance (**Figure 2D**). According to the Refseq annotation of the porcine genome, the segment of extended autozygosity encompasses 15 genes (**Figure 2E**).

We suspected that the mutation causing the sperm tail defect affects a transcript that is abundant in the testes. We found expression levels greater than 10 transcripts per million for four genes (*DNAH17*, *BIRC5*, *SYNGR2*, *SEC14L1*) that were annotated within the 1.11 Mb segment of extended homozygosity (**Figure 2E**). Among those, only *DNAH17* encoding the dynein axonemal heavy chain 17, shows a highly testis-biased expression in humans (https://gtexportal.org/home/), sheep (http://biogps.org/dataset/BDS_00015/sheep-atlas/) (Clark *et al.*, 2017) and cattle (http://cattlegeneatlas.roslin.ed.ac.uk/) (Fang *et al.*, 2020). Pathogenic alleles in human and murine orthologs of *DNAH17* have been detected in males that produce immotile sperm with multiple morphological abnormalities of the flagella (Whitfield *et al.*, 2019),(Zhang *et al.*, 2020). Thus, we considered *DNAH17* as a compelling positional and functional candidate gene for a sperm flagella defect in boars.

### An intronic 13-bp deletion is associated with the sperm tail defect

We sequenced five affected boars to an average coverage of 10.9-fold using short paired-end reads. We also considered 82 sequenced pigs from our in-house variant database that were not affected by the sperm tail disorder. The analysis of sequencing depth at the proximal region of SSC12 did not reveal large deletions or duplications that segregate with the haplotype. To identify candidate causal mutations, we considered 18,662 variants (14,806 SNPs and 3,856 Indels) that were detected within the 1.11 Mb interval (between 3,473,632 and 4,587,759 bp) of extended homozygosity. Of the 82 sequenced control animals, 65 also had array-called genotypes. Three of the 65 pigs that had array- and sequence-called genotypes carried the top haplotype in heterozygous state, yielding a haplotype frequency of 2% in the sequenced control animals.

Of 18,662 positional candidate causal variants, a single variant was compatible with recessive inheritance of the sperm tail defect: a 13-bp deletion at Chr12:3,556,401-3,556,414 bp (XM_021066525.1:c.8510-17_8510-5del) located in an intron of *DNAH17*. The 13-bp location resides within the top window from the haplotype-based association study. Sequence variant genotyping using *GATK* indicated that the five sequenced boars with the sperm tail defect were homozygous for the deletion. Only the three boars from the control group, that carried the top haplotype in the heterozygous state, also carried the 13-bp deletion in the heterozygous state. The deletion did not occur in the homozygous state in animals from the control group. According to the variant classification algorithm of the VEP software, the intronic 13-bp deletion was annotated as “splice_region_variant & intron_variant”. Its impact on protein function was predicted to be low.

### RNA sequencing reveals that the 13-bp deletion causes skipping of DNAH17 exon 56

Closer inspection of the 13-bp deletion revealed that it excises an intronic pyrimidine-rich sequence containing a continuous stretch of eight pyrimidine bases between 5 and 17 bases upstream of a canonical 3’ splice site of *DNAH17* exon 56 (**Figure 3A**). The VEP plugin SpliceRegion.pm indicated that the deletion coincides with a putative polypyrimidine tract. A branch point consensus sequence (yUnAy) (Gao *et al.*, 2008) was predicted (z-score: 4.79) 2-6 nucleotides upstream the 13-bp deletion. The pyrimidine content between the predicted branch point adenine and the splice acceptor site at the 3’ end of intron 55 is 68.4%. Considering that the polypyrimidine tract is an important *cis*-acting element for spliceosome assembly in canonical “GT-AG”-type acceptor splice sites (Coolidge *et al.*, 1997), we suspected that the 13-bp deletion perturbs pre-mRNA splicing. *In silico* splice site prediction using NNSPLICE indicated (prediction score 0.46) that the 13-bp deletion likely prevents recognition of the 3’ splice acceptor site at the intron-exon boundary of exon 56, thus possibly causing exon skipping.

**Figure 3:**
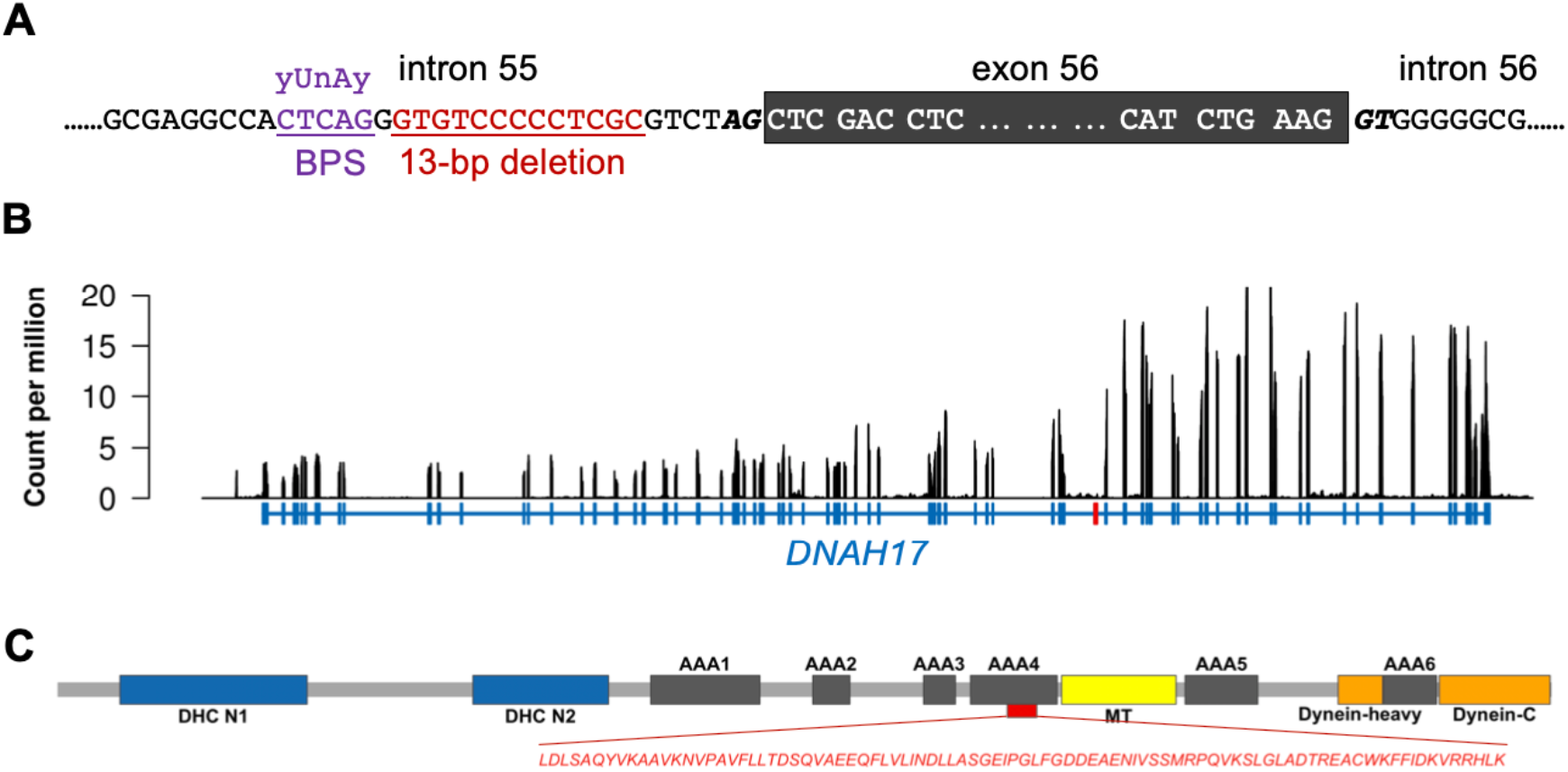
A 13-bp deletion perturbs splicing of *DNAH17*. Genomic sequence at the intron-exon boundaries of exon 56 of porcine *DNAH17* (XM_021066525.1) (A). The grey background indicates the sequence of exon 56. Red color highlights 13 nucleotides that are deleted in boars with the sperm tail defect. Splice acceptor and donor sites are represented in bold-italic type. Violet color indicates the predicted (z-score: 4.79) branch point sequence (BPS). Expression of DNAH17 (in counts per million) in testes of two Swiss Large White boars (SAMEA6832284, SAMEA6832285) homozygous for the 13-bp deletion (B). The number of reads covering a position was divided by the total number of reads (in million) mapped to transcripts, and averaged across both boars. Blue color indicates the exon-intron structure of porcine *DNAH17*. Exon 56 is displayed in red color. DNAH17 domains according to Uniprot (protein-ID: A0A287AFU3) (C). Different colored boxes represent conserved domains in the stem of dynein heavy chains (DHC), the microtubule-binding region (MT) and six ATPase domains (ATPases associated with diverse cellular activities, AAA1-AAA6). Skipping of exon 56 truncates 89 amino acids (red color) from AAA4.

Exon 56 of porcine *DNAH17* is regularly expressed in testis tissue of pubertal boars (**Supplemental file S5**). In order to examine if the loss of the polypyrimidine tract upstream exon 56 perturbs splicing, we sequenced RNA purified from testis tissue of two boars that produced defective sperm and were homozygous for the 13-bp deletion. We aligned 49 and 70 million paired-end (2×150 nt) RNA sequencing reads to the porcine transcriptome to quantify transcript and exon abundance. An average of 49 *DNAH17* transcripts per million (TPM) were detected in the two boars homozygous for the 13-bp deletion. However, no reads aligned to the sequence of exon 56 corroborating that the 13-bp deletion prevents recognition of the 3’ splice site, thereby causing exon skipping (**Figure 3B**).

The skipping of exon 56 causes an in-frame loss of 89 amino acids (residues 2836-2924) representing 2% of porcine DNAH17 (XP_020922184.1). DNAH17 contains six evolutionarily conserved “ATPase associated with a variety of cellular activities (AAA)”-domains that are linked together as an asymmetric hexameric ring required to power the beating movement of sperm flagella (Gleave *et al.*, 2014),(Snider *et al.*, 2008). The 89 amino acids are truncated from AAA4 of DNAH17 (**Figure 3C**).

## DISCUSSION

Eight Swiss Large White boars produced sperm with multiple morphological abnormalities of the flagella. Considering that all sperm were immotile, the boars were judged to be infertile and not used for breeding. Because a majority of the sperm were viable, fertilization might be possible using intracytoplasmic sperm injection (ICSI) (Nijs *et al.*, 1996),(Ortega *et al.*, 2011). While ICSI is not routinely applied in pig breeding programs, it could enable the reproduction of boars with sperm tail defects that prevent fertilization otherwise. However, success rates after ICSI may vary considerably in individuals with asthenoteratozoospermia (Wambergue *et al.*, 2016). Interestingly, ICSI success also varies in individuals with pathogenic *DNAH17* alleles (Whitfield *et al.*, 2019). In any case, the application of ICSI in livestock populations seems unwarranted if the underlying male-factor infertility follows recessive inheritance, because a defective allele would be transmitted to the offspring and might increase allele frequency in the population.

Apart from producing sperm with defective tails, the boars were healthy. Diseases and environmental stressors were less likely to cause the defective sperm flagella (Jurewicz *et al.*, 2009), because the boars were maintained at a semen collection center with many other boars that produced normal semen. Idiopathic infertility and poor semen quality in healthy males may result from pathogenic alleles in genes that are specifically expressed in the male reproductive tract (Pausch *et al.*, 2014),(Iso-Touru *et al.*, 2019),(Pausch *et al.*, 2016b),(Noskova *et al.*, 2020). The eight affected boars were inbred on a common ancestor supporting the hypothesis of a shared genetic etiology. While the presence of common ancestors in the pedigrees of all affected boars is compatible with recessive inheritance, it does not rule out other modes of inheritance (Bourneuf *et al.*, 2017). Considering that all affected but only few fertile boars were inbred on the common ancestor, a recessive mode of inheritance was likely. A mating between two heterozygous carriers resulted in the birth of eight female and three male piglets (5 wt/wt, 2 wt/mt, 4 mt/mt). The three male piglets were homozygous carriers of the 13-bp deletion and manifested the sperm tail defect, thus corroborating recessive inheritance.

The morphological and ultrastructural flagella defects of the eight Swiss Large White boars are similar to those observed in Finnish Yorkshire boars that are homozygous for a recessive loss-of-function allele in *KPL2* (a.k.a. *SPEF2*) (Andersson *et al.*, 2000),(Sironen *et al.*, 2006). Although cross-breeding between Finnish Yorkshire and Swiss Large White pigs had not been documented recently, the defective allele could be located on an ancient shared haplotype (Pausch *et al.*, 2016a),(Schwarzenbacher *et al.*, 2016). However, the genomic region on chromosome 16 encompassing *SPEF2* is not associated with the sperm defect of the Swiss Large White boars, indicating genetic heterogeneity.

SNPs on chromosomes 11, 12 and 13 were significantly associated with the sperm disorder using Fisher’s exact test of allelic association. This result was puzzling because an oligogenic inheritance of the sperm flagella defect was unlikely.

Phenotypic misclassification and genetic heterogeneity could lead to an inconclusive association study (Manchia *et al.*, 2013). However, the morphological abnormalities of the sperm flagella were strikingly similar in the ejaculates of eight boars. Moreover, pedigree analysis suggested monogenic recessive inheritance. Principal components analysis and a genomic inflation factor of 1.55 indicated that population stratification confounded our SNP-based association study (Price *et al.*, 2010). Although Fisher’s exact tests have been widely applied to map binary traits (Balding, 2006), these tests are prone to type I errors when cases and controls differ in their ancestries. A linear mixed model-based association study successfully controlled for population stratification and revealed two associated SNPs at SSC12. However, the SNPs were 6 million bp away from each other and the association signal was not very strong (P=1.2 × 10^−9^). Following the approach from (Hiltpold *et al.*, 2020), we fitted the top principal components of a genomic relationship matrix as covariates in a haplotype-based association model to take population stratification into account. A compelling association signal (P=4.22 × 10^−15^) at chromosome 12 remained while all other association signals disappeared. In agreement with previous studies (Price *et al.*, 2006),(Pausch *et al.*, 2011),(Hiltpold *et al.*, 2020) and evidenced by a low inflation factor, the principal components-based correction successfully eliminated spurious signals that arose due to population stratification. Considering that the most significantly associated haplotype also encompasses the causal variant for the sperm flagella defect, our findings corroborate that genome-wide haplotype-based association testing offers a powerful approach to identify trait-associated regions in stratified mapping cohorts (Kadri *et al.*, 2014),(Hiltpold *et al.*, 2020),(Pausch *et al.*, 2016a).

The haplotype associated with multiple morphological abnormalities of the sperm flagella carries a 13-bp deletion in an intron of *DNAH17* encoding dynein axonemal heavy chain 17. Our results show that DNAH17 is abundant in porcine testis tissue. Dynein axonemal heavy chains are required for the axonemal assembly and the beating movement of the flagella (Inaba, 2003),(Ben Khelifa *et al.*, 2014),(Tu *et al.*, 2019),(Sironen *et al.*, 2020),(Liu *et al.*, 2020),(Whitfield *et al.*, 2019),(Zhang *et al.*, 2020),(Sha *et al.*, 2020). Pathogenic alleles in human and murine *DNAH17* manifest in sperm with an abnormal mitochondrial sheath and cytoplasmic droplets, as well as axonemal disorganization including absence of the central pair and missing peripheral microtubule doublets (Whitfield *et al.*, 2019), (Zhang *et al.*, 2020),*(Sha et al.*, 2020). Boars that are homozygous for a 13-bp deletion in intron 55 of *DNAH17* also produce sperm with multiple morphological and ultrastructural abnormalities of the flagella. Although annotated as a low impact variant, the intronic 13-bp deletion perturbs pre-mRNA splicing because it disrupts the polypyrimidine tract in intron 55 of *DNAH17*. The mutant DNAH17 lacks part of the AAA4 domain which likely compromises axonemal assembly and the beating movement of the sperm flagellum in the homozygous state. The assembly of the flagellar axoneme is less disorganized when other domains of DNAH17 are affected by pathogenic variants (Whitfield *et al.*, 2019),(Zhang *et al.*, 2020),(Sha *et al.*, 2020), likely because an intact AAA4 domain is essential to physiological DNAH17 function (Snider *et al.*, 2008). Thus, our findings suggest that the intronic 13-bp deletion is a loss-of-function allele.

We previously identified mutations at splice donor sites and nearby splice acceptor sites that manifest in male reproductive disorders due to aberrant splicing (Iso-Touru *et al.*, 2019),(Hiltpold *et al.*, 2020). Our present study reveals an intronic deletion from 5 to 17 nucleotides upstream a canonical 3’ splice site that perturbs splicing as the most likely causal variant for a novel porcine sperm flagella defect. The 13-bp deletion compromises splicing because it excises a polypyrimidine tract upstream *DNAH17* exon 56. Phenotypic consequences arising from polypyrimidine tract mutations had rarely been described so far (Lefévre *et al.*, 2002),(Sartelet *et al.*, 2015),(Abramowicz and Gos, 2018). To the best of our knowledge, our study reports the first phenotype-genotype association for a mutation affecting an intronic polypyrimidine tract in pigs. We provide evidence that the loss of the polypyrimidine tract in porcine *DNAH17* intron 55 prevents recognition of the 3’ splice site which leads to the skipping of exon 56, thus causing defective sperm flagella. Sequence motifs that govern the assembly of the spliceosome may be more distant to 3’ splice sites than the polypyrimidine tract in porcine *DNAH17* intron 55 (Zhang *et al.*, 2017). However, standard sequence variant annotation tools are largely blind to putative consequences arising from mutations within intronic motifs. Thus, the systematic characterization of branch points and polypyrimidine tracts seems warranted to refine the functional classification of intronic variants.

Our findings enable the monitoring of the 13-bp deletion and unambiguous identification of carrier and homozygous animals using direct gene testing. While homozygosity for the 13-bp deletion is easily recognized in artificial insemination boars using microscopic semen analysis, it may remain undetected in sows and natural service boars. Affected natural service boars will be noticed after few matings due to low fertility. In agreement with previous findings on loss-of-function alleles in human and murine *DNAH17* (Whitfield *et al.*, 2019),(Zhang *et al.*, 2020),(Sha *et al.*, 2020), homozygous mutation carriers were healthy. Considering that the frequency of the 13-bp deletion is low and the majority of sows is inseminated artificially, economic losses due to unsuccessful breeding with homozygous boars are negligible in the Swiss Large White population.

The number of recessive conditions detected in livestock populations increases steadily (Nicholas and Hobbs, 2014). Low effective population size, founder effects and inbreeding favor the manifestation of recessive conditions. Recent advances in genome-wide genotyping and sequencing offer powerful tools for rapid detection of causal variants underpinning inherited conditions (Bourneuf *et al.*, 2017). The phenotypic and economic consequences of the 13-bp deletion in *DNAH17* are less detrimental than mutations that manifest in malformed and non-viable piglets (Fang *et al.*, 2020),(Derks *et al.*, 2018). However, alleles that compromise semen quality and male fertility in otherwise healthy individuals may attain high frequency in the absence of deleterious manifestations in females (Pausch *et al.*, 2014),(Hiltpold *et al.*, 2020). The 13-bp deletion in *DNAH17* adds to a catalogue of sequence variants that cause male-factor infertility in pigs (Sironen *et al.*, 2011),(Sironen *et al.*, 2006),(Noskova *et al.*, 2020). It remains an open question how an increasing number of undesired recessive alleles may be considered appropriately in livestock populations (Cole, 2015),(Upperman *et al.*, 2019). Although the newly identified 13-bp deletion does neither compromise the welfare of homozygous animals nor result in huge economic losses, it’s frequency should be kept at a low level to prevent the birth of homozygous boars that manifest a sterilizing sperm tail disorder.

## ACKNOWLEDGEMENTS

We acknowledge financial support from SUISAG, Micarna SA and the ETH Zürich Foundation. We thank Dr. Cecilia Bebeacua (ScopeM) for support in transmission electron microscopy. We are thankful for the excellent technical support provided by the ETH Zürich technology platforms FGCZ (https://fgcz.ch/), ScopeM (https://scopem.ethz.ch/), and Genetic Diversity Centre (https://gdc.ethz.ch/), respectively, for sequencing, microscopy and DNA fragment analysis.

## SUPPLEMENTAL FILES

### Supplemental file 1

Title: Accession numbers of 87 boars

Format: xlsx

Description: Accession numbers of the sequence data of five affected (asthenozoospermic) and 82 unaffected (control) boars. The third column indicates the haplotype status (0 – not carrier, 1 – heterozygous carrier, 2 – homozygous carrier) of 65 sequenced boars that also had Illumina PorcineSNP60 BeadChip-called genotypes.

### Supplemental file 2

Format: jpg

Title: Sperm viability assessment by eosin-nigrosin staining

Description: Representative light microscopic image of sperm from an affected boar stained with eosin-nigrosin. The sperm were flushed post mortem from the epididymis of Boar_1249. Live sperm appear white in color. Dead sperm appear pink in color because they take up eosin due to loss of membrane integrity.

### Supplemental file 3

Title: Disorganized flagellar axonemes in two affected boars

Format: png

Description: Cross-sections of sperm from two affected boars reveal multiple morphological and ultrastructural abnormalities of the flagella including the absence of outer or inner doublets, secondary flagella-like structures within the same membrane, and flagellar components in the cytoplasmic droplet-like structure.

### Supplemental file 4

Title: Principal components analysis of cases and controls

Format: png

Description: Scatter plot of the top five eigenvectors of a genomic relationship matrix. Grey and red color indicates 100 fertile and six asthenoteratozoospermic boars from the Swiss Large White population. The raw data underlying the plot (genotypes, haplotypes, eigenvectors) are available at available at https://doi.org/10.5281/zenodo.4081475.

### Supplemental file 5

Title: Expression profile of porcine *DNAH17* in pubertal boars

Format: png

Description: Expression of *DNAH17* quantified using testis RNAseq alignments of four pubertal boars (SRR2564762, SRR8237161, SRR8237162, SRR8237165). The number of reads covering a position was extracted from coordinate sorted BAM files using the MOSDEPTH software, divided by the total number of reads mapped to transcripts. Blue color indicates the exon-intron structure of porcine *DNAH17*. Exon 56 is displayed in red color. Although exon expression is somewhat variable between the samples, the expression of exon 56 is evident in all samples. Raw data underlying the plot are available at the European Nucleotide Archive (ENA) of the EMBL at the BioProject PRJNA506525.

