## Supplementary figures and images for "Infertility due to defective sperm flagella caused by an intronic deletion in DNAH17 that perturbs splicing"

### Supplemental_file_2.jpg

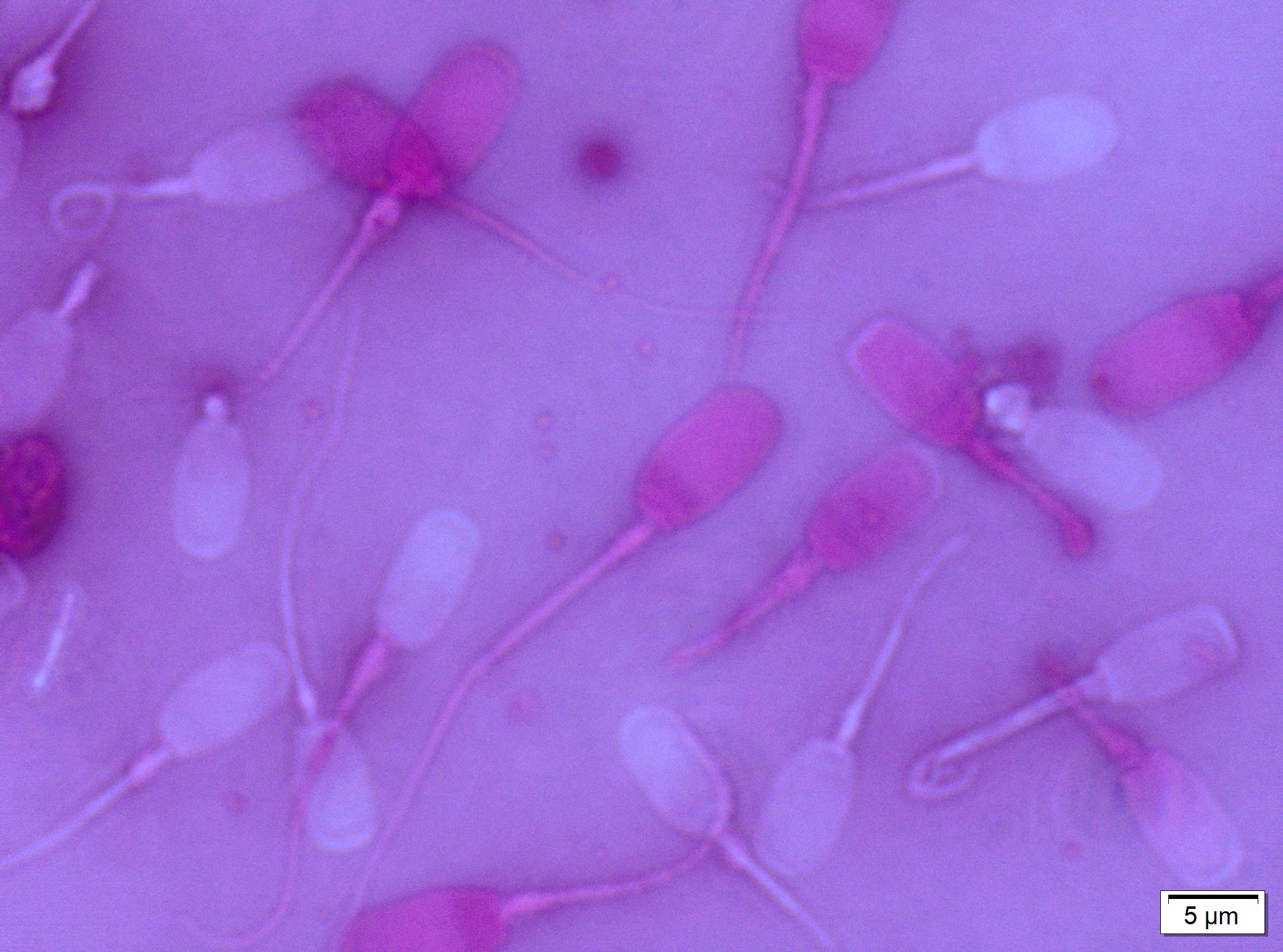

### Supplemental_file_3.png

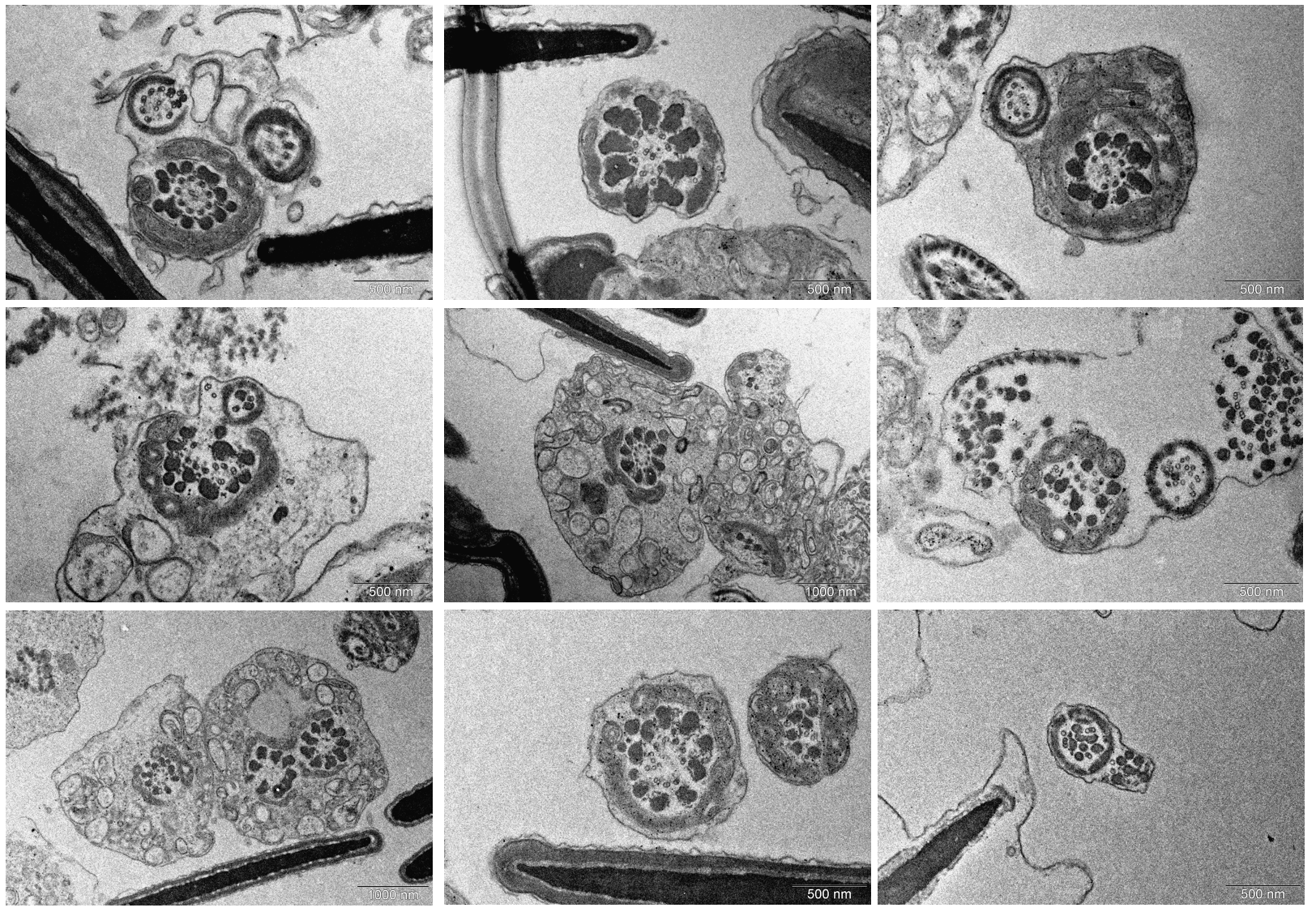

### Supplemental_file_4.png

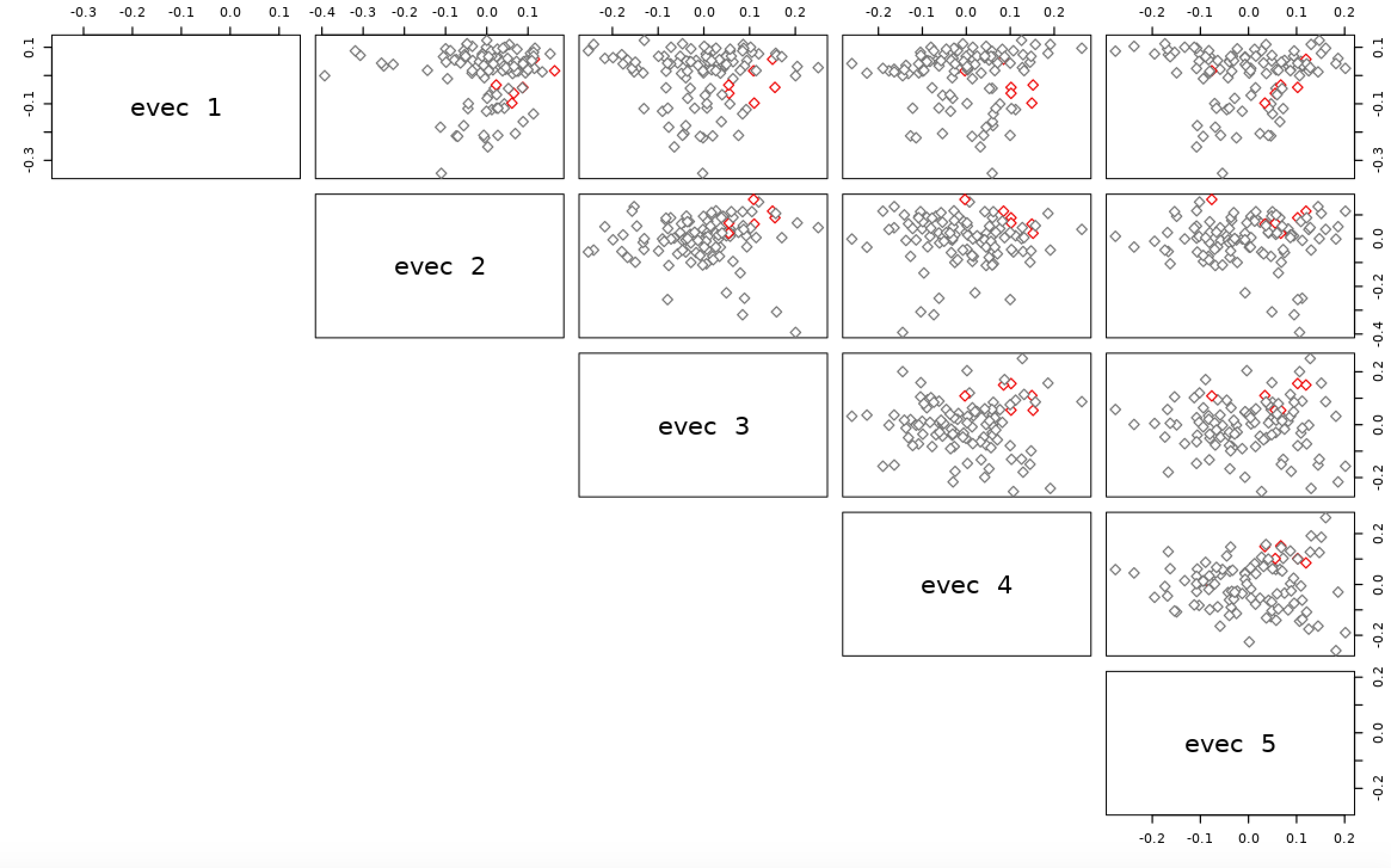

### Supplemental_file_5.png

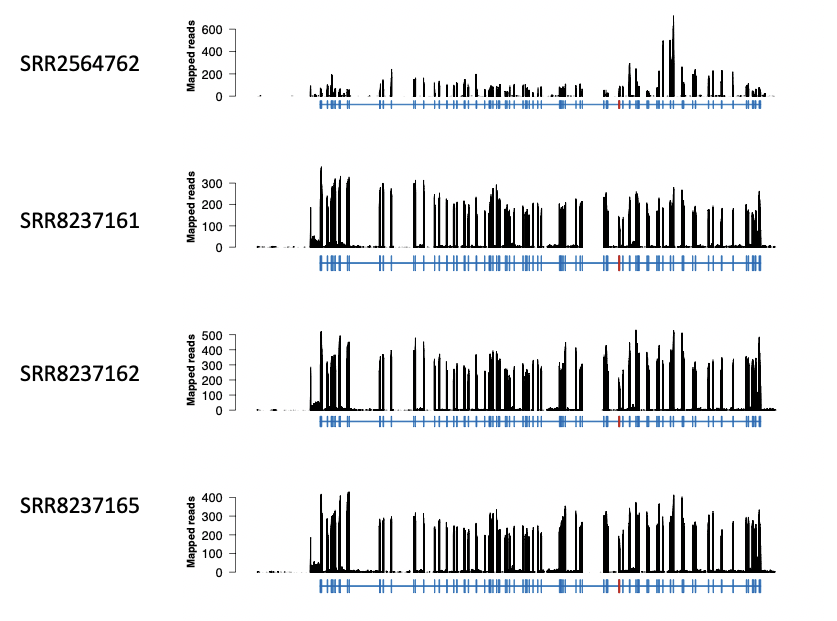
